# Whole-Brain opto-fMRI Map of Mouse VTA Dopaminergic Activation Reflects Structural Projections with Small but Significant Deviations

**DOI:** 10.1101/2020.04.03.023648

**Authors:** Horea-Ioan Ioanas, Bechara John Saab, Markus Rudin

## Abstract

Ascending dopaminergic projections from neurons located in the Ventral Tegmental Area (VTA) are key to the etiology, dysfunction, and control of motivation, learning, and addiction. Due to evolutionary conservation of this nucleus and the extensive use of mice as disease models, establishing an assay for VTA dopaminergic signalling in the mouse brain is crucial for the translational investigation of motivational control as well as of neuronal function phenotypes for diseases and interventions. In this article we use optogenetic stimulation directed at VTA dopaminergic neurons in combination with functional Magnetic Resonance Imaging (fMRI), a method widely used in human deep brain imaging. We present a comprehensive assay producing the first whole-brain opto-fMRI map of dopaminergic activation in the mouse, and show that VTA dopaminergic system function is consistent with its structural VTA projections, diverging only in a few key aspects. While the activation map predominantly highlights target areas according to their relative projection densities (e.g. strong activation of the nucleus accumbens and low activation of the hippocampus), it also includes areas for which a structural connection is not well established (such as the dorsomedial striatum). We further detail the variability of the assay with regard to multiple experimental parameters, including stimulation protocol and implant position, and provide evidence-based recommendations for assay reuse, publishing both reference results and a reference analysis workflow implementation.

## Background

The dopaminergic system consists of a strongly localized, and widely projecting set of neurons with cell bodies clustered in the midbrain into two lateralized nucleus pairs, the Substantia Nigra pars compacta (SNc) and the Ventral Tegmental Area (VTA, fig. 1a). On account of the small number of dopaminergic neurons (≈ 300, 000 in humans [1], ≈ 10, 000 in rats [2], and ≈ 4, 000 in mice [3]), tractography commonly fails to resolve the degree centrality of this neurotransmitter system, precluding it from being a prominent node in such graph representations of the brain. However, it is precisely the small number of widely branching and similar neurons, which makes the dopaminergic system a credible candidate for truly node-like function in coordinating brain activity. As is expected given such salient features, the system is widely implicated in neuropsychiatric phenomena (including addiction [4, 5], attentional control [6], motivation [7], creativity [8], personality [9], neurodegeneration [10], and schizophrenia [11]), and is a common target for pharmacological interventions. Lastly, due to high evolutionary conservation [12], the dopaminergic system is also an excellent candidate for translational study.

**Figure 1:**
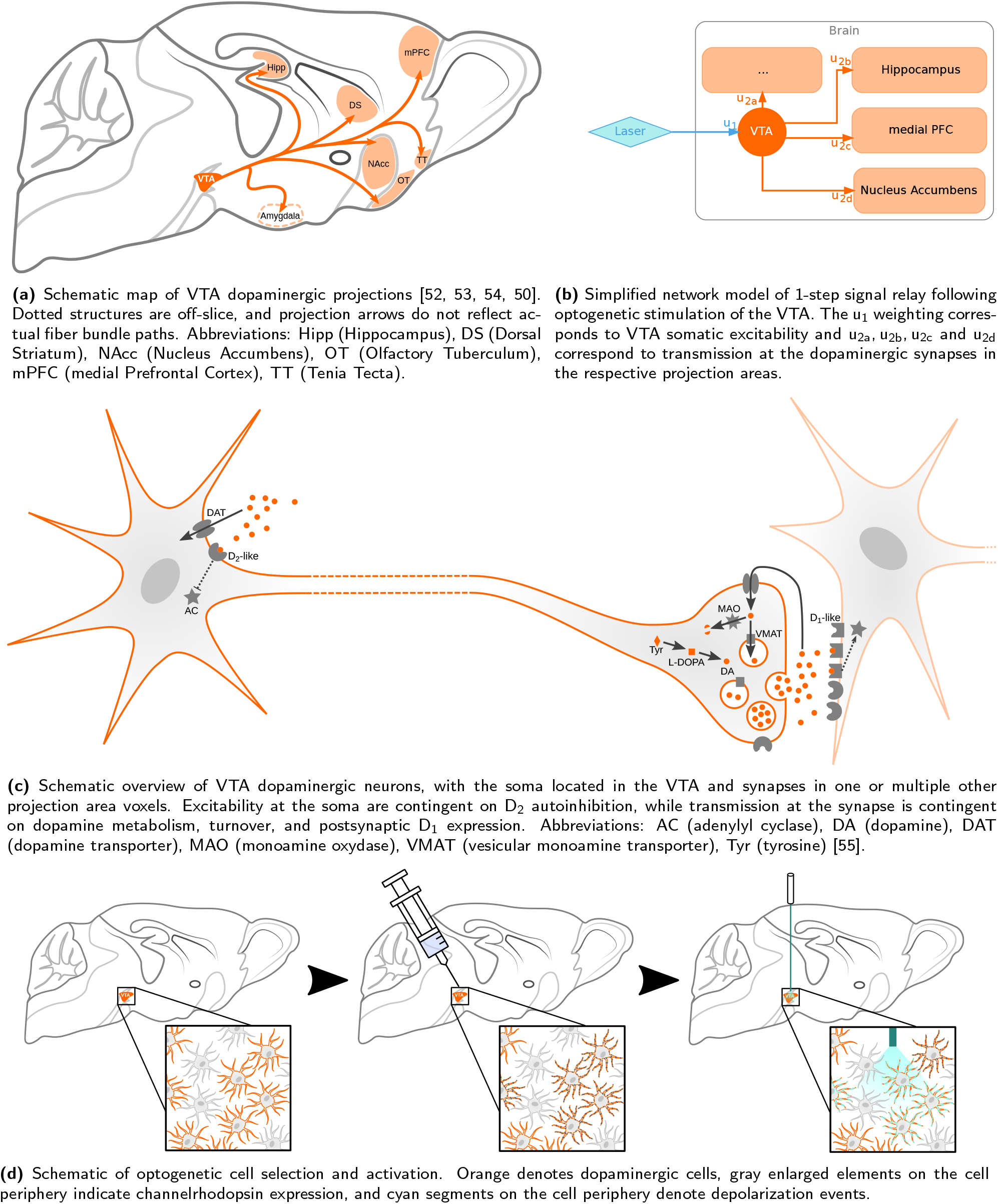
The cell biological compartmentalization of dopaminergic neurotransmission (and susceptibility to psychopharmacology) can partly be mapped onto neuroanatomical features by a simple network model, using optogenetics. Depicted are schematic overviews of the VTA dopaminergic system at various spatial resolutions.

**Figure 2:**
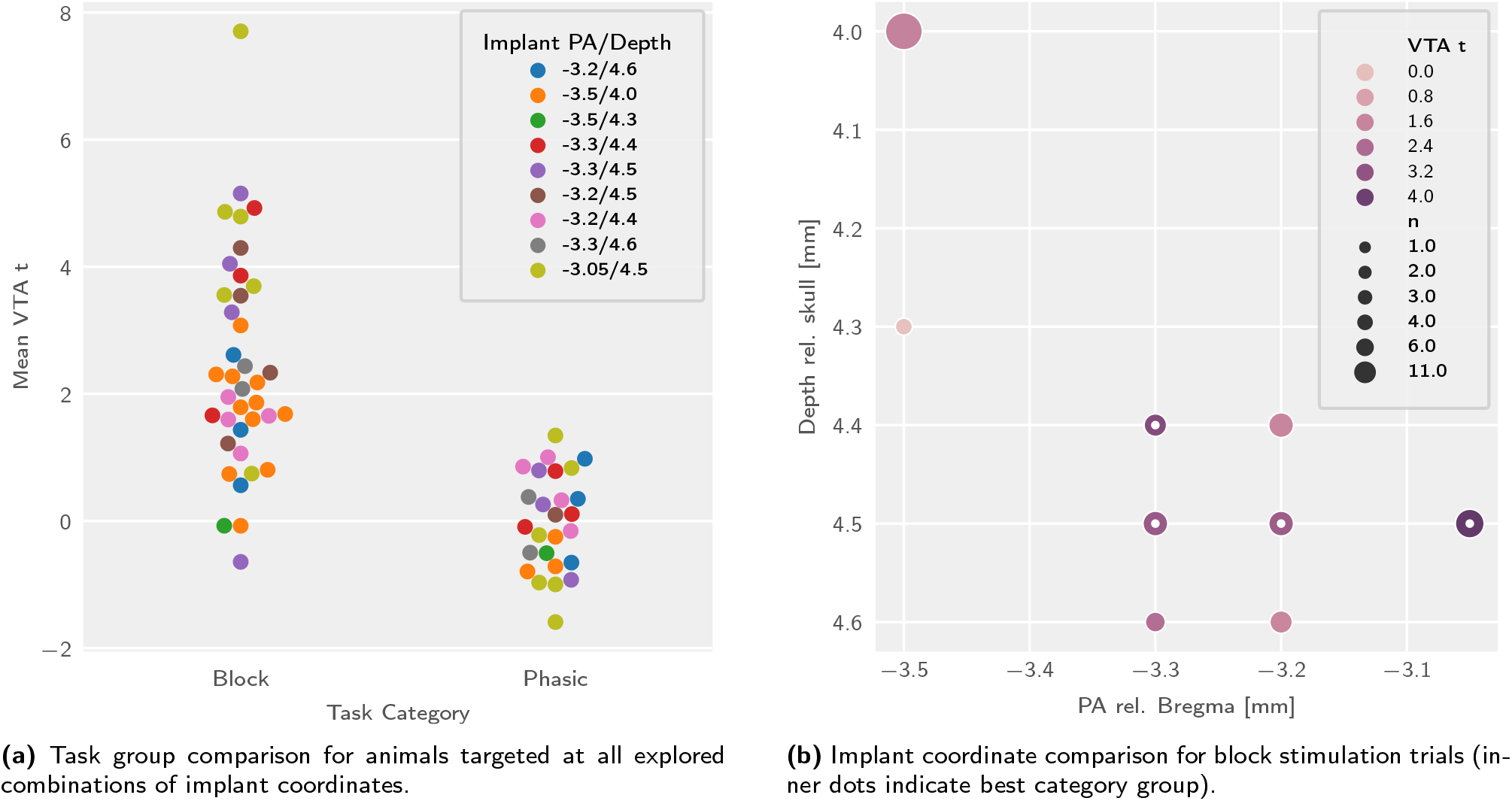
VTA activation is sensitive to the stimulation protocol category and the implant coordinates, with different trends in block and phasic stimulation trials. Depicted are multifactorial (protocol and implant coordinates) comparisons of signal intensity in the VTA region of interest. Abbreviations: n (sample size), PA (posteroanterior), rel. (relative to).

Imaging a neurotransmitter system comprised of a small number of cells based only on spontaneous activity is highly unreliable due to an intrinsically low signal to noise ratio (SNR). This limitation can, however, be overcome by introducing exogenous stimulation. While the colocalization of widely projecting dopaminergic cell bodies into nuclei renders temporally precise and population-wide targeting feasible, dopaminergic nuclei also contain notable sub-populations of non-dopaminergic cells, which may confound an intended dopaminergic read-out [13]. In order to specifically target dopaminergic cells, they need to be sensitized to an otherwise inert stimulus in a transcription-dependent manner. This can be achieved via optogenetics, which is based on light-stimulation of cells expressing light-sensitive proteins such as channelrhodopsin [14]. Cell-type selectivity can be achieved by Cre-conditional channelrhodopsin vector delivery [15] to transgenic animals expressing Cre-recombinase under a dopaminergic promoter. Following protein expression, stimuli can be delivered via an implanted optic fiber. The combination of this stimulation method with fMRI is commonly referred to as opto-fMRI and can provide information on functional connectivity between a primary activation site and associated projection areas [16, 17].

Key questions surrounding VTA function in pre-clinical models are, firstly, method feasibility in animal models more accessible to transgenic techniques, such as the mouse; and secondly, a mapping of the efferent spectrum for dopaminergic VTA output. In particular, in the study of the Rat VTA, it has both been suggested that the efferent dopaminergic spectrum encompasses but extends beyond well-documented structural projections [18] — or alternatively, that VTA dopaminergic efferences are comparatively sparse and that based on translational insight the dopaminergic paradigm of motivation-related VTA function could be questioned [19].

The current study of whole-brain VTA dopaminergic function in mice aims to produce three novel research outputs. Firstly, a proof-of-principle documenting the feasibility of midbrain dopaminergic opto-fMRI in the mouse should be demonstrated, using a protocol which affords qualitative comparability with extant rat data, such as block stimulation and right VTA targeting. Pursuing open questions in the field, results should be quantitatively benchmarked with respect to histologically documented structural projections in the mouse. Secondly, the procedure needs to be optimized by systematic variation of experimental parameters (such as targeting and stimulation protocol variations) in order to ascertain reliability and reproducibility, as is required for a general-purpose dopaminergic system assay. Lastly, a reference neurophenotype of stimulus-evoked dopaminergic function (represented as a brain-wide voxelwise map) should be published in standard space to facilitate co-registered data integration, operative targeting, and comparative evaluation of pathology or treatment induced effects.

These goals presuppose not only the production of experimental data, but also the development of a transparent, reliable, and publicly accessible analysis workflow, which leverages pre-existing standards for mouse brain data processing [20] and extends them to the statistical analysis.

## Methods

### Animal Preparation

VTA dopaminergic neurons were specifically targeted via optogenetic stimulation. As shown in fig. 1d, this entails a triple selection process. Firstly, cells are selected based on gene expression (via a transgenic mouse strain), secondly the location is selected based on the injection site, and thirdly, activation is based on the overlap of the aforementioned selection steps with the irradiation volume covered by the optic fiber.

A C57BL/6-based mouse strain was chosen, which expresses Cre recombinase under the dopamine transporter (DAT) promoter [21]. Transgenic construct presence was assessed via polymerase chain reaction (PCR) for the Cre construct, using the forward primer ACCAGCCAGCTATCAACTCG and the reverse primer TTGCCCCTGTTTCACTATCC. A total of 25 transgenic animals and 7 wild type control animals are included in the study. The animal sample consisted of 18 males and 14 females, with a group average age of 302 days (standard deviation 143 days) at the study onset. The sample size was determined based on the range found sufficient to uncover opto-fMRI results in the mouse serotonergic system [17].

The right VTA (fig. 3e, green contour) of the animals was injected with a recombinant Adeno-Associated Virus (rAAV) solution. The vector delivered a plasmid containing a floxed channelrhodopsin and YFP construct: pAAV-EF1a-double floxed-hChR2(H134R)-EYFP-WPRE-HGHpA, gifted to a public repository by Karl Deisseroth (Addgene plasmid #20298). Viral vectors and plasmids were produced by the Viral Vector Facility (VVF) of the Neuroscience Center Zurich (Zentrum für Neurowissenschaften Zürich, ZNZ). The solution was prepared at a titer of 5.7 × 10^12^ vg/ml and volumes from 0.8 to 1.6 μl were injected into the right VTA. Injection coordinates ranged in the posteroanterior (PA) direction from −3.5 to −3.05 mm (relative to bregma), in depth from 4.0 to 4.4 mm (relative to the skull), and were located 0.5 mm right of the midline. Construct expression was ascertained post mortem by fluorescent microscopy of formaldehyde fixed 200 μm brain slices.

**Figure 3:**
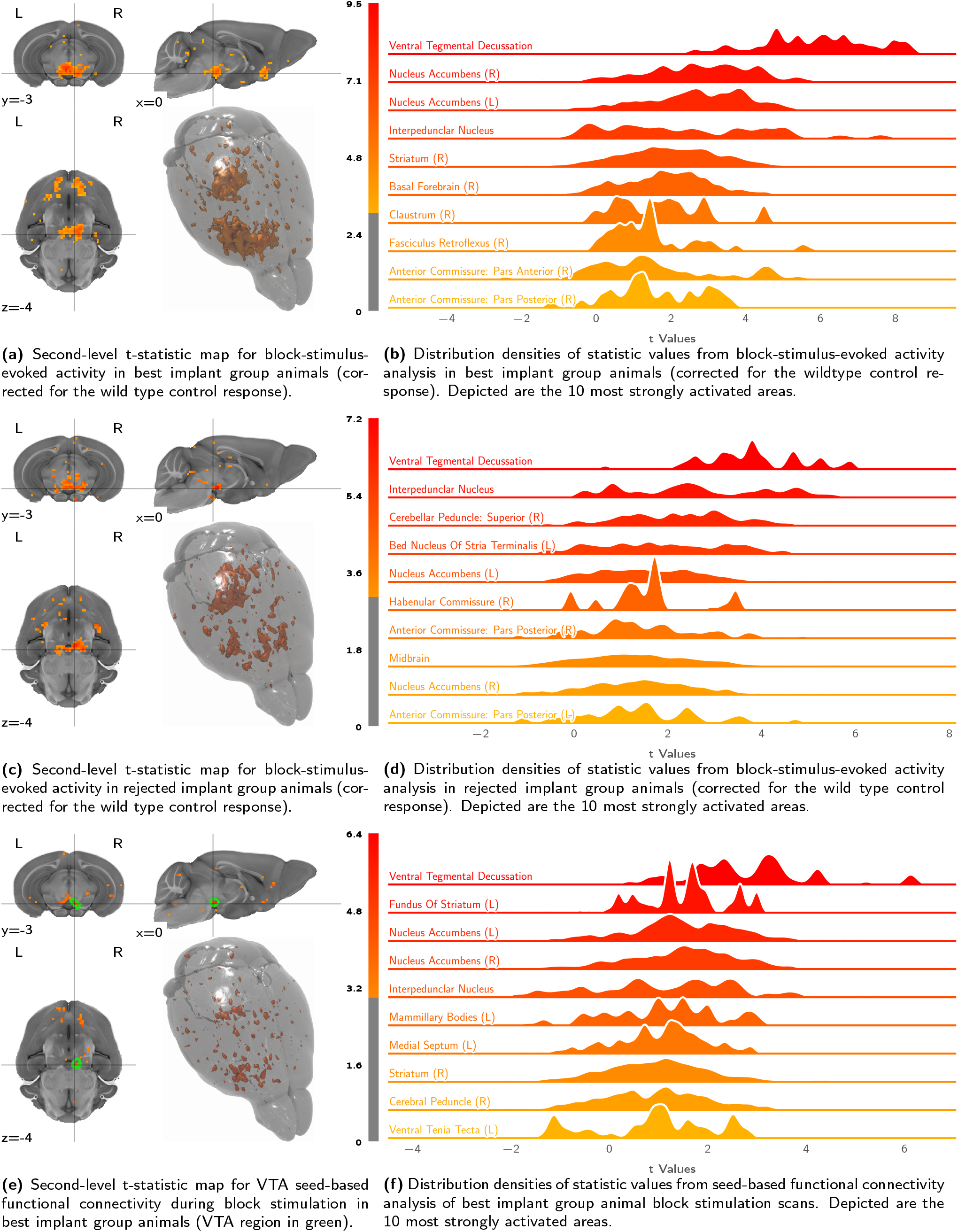
Block stimulation elicits strong ventral striatal activity in the best implant group, more rostrally weighted activity in the rejected implant group, and generates similar but weaker contrasts for VTA seed-based analysis. The figures show volumetric population t-statistic maps **(a**, **e**, **c)** thresholded at t ≥ 3 and centered on the VTA target, as well as a break-down of activation along atlas parcellation regions **(b**, **d**, **f)**.

For optical stimulation, animals were fitted with an optic fiber implant (l = 4.7 mm d = 400 μm NA = 0.22) targeting the right VTA, at least two weeks before imaging. Implant target coordinates ranged in the PA direction from −3.5 to −3.05 mm (relative to bregma), in depth from 4.0 to 4.6 mm (relative to the skull), and were located 0.5 to 0.55 mm right of the midline.

Stimulation was delivered via an Omicron LuxX 488-60 laser (488 nm), tuned to a power of 30 mW at contact with the fiber implant, according to the protocols listed in tables S1 to S7. Stimulation protocols were delivered to the laser and recorded to disk via the COSplayer device [22]. Animal physiology, preparation, and measurement metadata were tracked with the LabbookDB database framework [23].

### MR Acquisition

Over the course of preparation and measurement, animals were provided a constant flow of air with an additional 20 % O_2_ gas (yielding a total O_2_ concentration of ≈ 36 %). For animal preparation, anesthesia was induced with 3 % isoflurane, and maintained at 2 to 3 % during preparation — contingent on animal reflexes. Animals were fixed to a heated MRI-compatible cradle via ear bars and a face mask equipped with a bite hook. A subcutaneous (s.c.; right dorsal) and intravenous (i.v.; tail vein) infusion line were applied. After animal fixation, a bolus of medetomidine hydrochloride (Domitor, Pfizer Pharmaceuticals, UK) was delivered s.c. to a total dose of 100 ng/(g BW) and the inhalation anesthetic was reduced to 1.5 % isoflurane. After a 5 min interval, the inhalation anesthetic was set to 0.5 % and medetomidine was continuously delivered at 200 ng/(g BW h) for the duration of the experiment. This anesthetic protocol is closely based on extensive research into animal preparation for fMRI [24].

All data were acquired with a Bruker Biospec system (7 T, 16 cm bore), and an in-house built transmit/receive surface coil, engineered to permit optic fiber implant protrusion.

Anatomical scans were acquired via a TurboRARE sequence, with a RARE factor of 8, an echo time (TE) of 30 ms, an inter-echo spacing of 10 ms, and a repetition time (TR) of 2.95 s. Thirty adjacent (no slice gap) coronal slices were recorded with a nominal inplane resolution of Δx(*ν*) = Δy(*ϕ*) = 75 μm (sampled as 180 voxels sagittally and 120 voxels horizontally), and a slice thickness of Δz(t) = 450 μm.

Functional scans were acquired with a gradient-echo EPI sequence, a flip angle of 60°, and TR/TE = 1000 ms/5.9 ms. Thirty adjacent (no slice gap) coronal slices were recorded with a nominal in-plane resolution of Δx(*ν*) = Δy(*ϕ*) = 225 μm (sampled as 60 voxels sagittally and 29 voxels horizontally), and a slice thickness of Δz(t) = 450 μm. Functional scans were acquired over a period of 25 min, totalling 1500 repetitions. Changes in cerebral blood volume (CBV) are measured as a proxy of neuronal activity following the administration of an intravascular iron oxide nanoparticle based contrast agent (Endorem, Laboratoire Guebet SA, France) [25]. The contrast agent (30.24 μg/(g BW)) is delivered as an i.v. bolus 10 min prior to the fMRI data acquisition, to achieve a pseudo steady-state blood concentration. This contrast is chosen to enable short echo-time imaging thereby minimizing artefacts caused by gradients in magnetic susceptibility.

The total duration of the scan session, including induction, preparation, and scanning (including the 10 min delay after contrast agent administration, taking place between the structural and functional scan) was approximately 80 min.

MR acquisition was performed blindly with respect to the implant parameter variation, the measurement order was not systematically separated between the conditions. All animal experiments and handling were performed in accordance with the relevant requirements of the Cantonal Veterinary Office of Zurich, under licence ZH263/14 and extension ZH128/18.

### Preprocessing

Data conversion from the proprietary ParaVision format was performed via the Bruker-to-BIDS repositing pipeline [26] of the SAMRI package (version `0.4` [27]). Following conversion, data was dummy-scan corrected, registered, and subject to controlled smoothing via the SAMRI Generic registration workflow [20]. As part of this processing, the first 10 volumes were discarded (automatically accounting for volumes excluded by the scanner software). Registration was performed using the standard SAMRI mouse-brain-optimized parameter set for ANTs [28] (version `2.3.1`). Data was transformed to a stereotactically oriented standard space (the DSURQEC template space, as distributed in the Mouse Brain Atlases Package [29], version `0.5.3`), which is based on a high-resolution T_2_-weighted atlas [30]. Controlled spatial smoothing was applied in the coronal plane up to 250 μm via the AFNI package [31] (version `19.1.05`).

The registered time course data was frequency filtered depending on the analysis workflow. For stimulus-evoked activity, the data was low-pass filtered at a period threshold of 225 s, and for seed-based functional connectivity, the data was band-pass filtered within a period range of 2 to 225 s.

### Statistics and Data

Volumetric data was modelled using functions from the FSL software package [32] (version `5.0.11`). First-level regression was applied to the temporally resolved volumetric data via FSL’s `glm` function, whereas the second-level analysis was applied to the first-level contrast and variance estimates via FSL’s `flameo`.

Stimulus-evoked first-level regression was performed using a convolution of the stimulus sequence with an opto-fMRI impulse response function, estimated by a beta fit of previously reported mouse opto-fMRI responses [17]. Seed-based functional connectivity analysis was performed by regressing the time course of the voxel most sensitive to the stimulus-evoked activity (per scan) in the VTA region of interest.

Brain parcellation for region-based evaluation was performed using a non-overlapping multi-center labelling [30, 33, 34, 35], as distributed in version `0.5.3` of the Mouse Brain Atlases data package [29]. The mapping operations were performed by a SAMRI function, using the nibabel [36] and nilearn [37] libraries (versions `2.3.1` and `0.5.0`, respectively). Classification of implant coordinates into “best” and “rejected” categories was performed via 1D k-means clustering, implemented in the scikit-learn library [38] (version `0.20.3`). Distribution density visualizations were created using the Scott bandwidth density estimator [39], as implemented in the seaborn software package (`0.9.0`).

Higher-level statistical modelling was performed with the Statsmodels software package [40] (version `0.9.9`), and the SciPy software package [41] (version `1.1.0`). Model parameters were estimated using the ordinary least squares method, and a type 3 analysis of variance (ANOVA) and a heteroscedasticity consistent covariance matrix [42] were employed to control estimate variability for unbalanced categories. All t-tests producing explicitly noted p-values are two-tailed.

The VTA structural projection data used to compare and contrast the activation maps produced in this study was sourced from the Allen Brain Institute (ABI) mouse brain connectome dataset [43]. As the target promoter of this study (DAT) is not included in the ABI connectome study, all available promoters were used (Sty17, Erbb4, Slc6a3, Th, Cck, Pdzk1ip1, Chrna2, Hdc, Slc18a2, Calb2, and Rasgrf2). Datasets with left-handed VTA injection sides were flipped to provide right-hand VTA projection estimates. The data was converted and registered to the DSURQEC template space by the ABI Connectivity Data Generator package [44]. For the second-level statistical comparison between functional activation and structural projection, individual activation (betas) and projection maps were normalized to a common scale by subtracting the average and dividing by the standard deviation.

Software management relevant for the exact reproduction of the aforementioned environment was performed via neuroscience package install instructions for the Gentoo Linux distribution [45].

All data analysis was performed on the entire dataset, without any data being removed, and in the absence of individual category investigation.

### Reproducibility and Open Data

The resulting t-statistic maps (i.e. the top-level data visualized in this document), which document the opto-fMRI dopaminergic map in the mouse model, are distributed along the source-code of all analyses [46]. The BIDS [47] data archive which serves as the raw data recourse for this document is openly distributed [48], as is the full instruction set for recreating this document from the aforementioned raw data [46]. The source code for this document and all data analysis shown herein is structured according to the RepSeP specifications [49].

## Results

Opto-fMRI experiments were carried out in C57BL/6 mice expressing Cre recombinase under the dopamine transporter promoter [21], with Cre-conditional viral vector induced expression of channelrhodopsin (ChR2) and yellow fluorescent protein (YFP) in the dopaminergic midbrain. Light stimuli were delivered via an optic fiber pointing above the right VTA. Different stimulation protocols were applied to the animals, consisting of variations within two main categories: block stimulation (with light stimuli delivered in continuous blocks of at least 8 s — tables S1 to S5) and phasic stimulation (with light stimuli delivered in short bursts of up to 1 s in lenght — tables S6 and S7). Additionally, the dataset details the effects of variation in the posteroainerior (PA) coordinates and the implant depth (equivalent to the dorsoventral coordinate of the fiber endpoint), specified relative to bregma and the skull surface, respectively.

In the analysis of the resulting data, the mean t-statistic for the stimulation regressor fit across the VTA region of interest is found sensitive to the stimulation protocol category (*F*_1,54_ = 47.26, *p* = 6.57 × 10^−9^), the stimulation target depth (*F*_4,54_ = 2.656, *p* = 0.043), the stimulation target PA coordinates (*F*_3,54_ = 3.063, *p* = 0.036), but not the interaction of the depth and PA target coordinates (*F*_12,54_ = 1.591, *p* = 0.16).

The break-up by phasic and block stimulation is shown in fig. 2 and significance is evaluated accounting for the entire statistical model, consisting of categorical terms for both the stimulus category and the coordinates. The phasic and block levels of the stimulation variable yield p-values of 0.063 and 1.87 × 10^−5^, respectively. Upon investigation of the t-statistic map, phasic stimulation further reveals no coherent activation pattern at the whole-brain level(fig. S3b).

The main and interaction effects of the implant coordinate variables are better described categorically than linearly (figs. S1 and 2b). Consequently, the most suitable implant coordinate group for the assay can best be determined on the basis of categorical classification of implant coordinates. We classify the implant coordinates into a “best” and a “rejected” group by k-means clustering the aggregate VTA t-statistic scores into two clusters, and find spatial coherence for the “best” coordinate group (categorization highlighted in fig. 2b).

For block stimulation, the best implant category group (fig. 3a) and the rejected implant category group (fig. 3c) show not only a difference in overall stimulus-evoked signal intensity, but also a difference in efferent distribution, with the rejected implant category efferent spectrum more strongly weighted towards caudal brain areas. This distinction specifically arises for implant categorization based on block scan VTA t-statistic means, and is not as salient if implants are categorized based on a posteroanterior implant coordinate delimiter (fig. S2).

The activation pattern elicited by block stimulation in the best implant category group shows strong coherent clusters of activation. The top activation areas are predominantly located in the right hemisphere, with highly significant laterality (*p* = 8.27 × 10^−7^) seen in the comparison of left and right hemisphere atlas parcellation region averages. Activation is seen in regions surrounding the stimulation site, such as the ventral tegmental decussation and the interpeduncular nucleus. The largest activation cluster encompasses well-known dopaminergic VTA projection areas in the subcortical rostroventral regions of the brain (nucleus accumbens, striatum, and the basal forebrain), with weaker activation observed in smaller structures in the vicinity of these regions, such as the fasciculus retroflexus, anterior commissure and the claustrum.

This activation pattern is is largely consistent with structural projection data, as published by the Allen Brain Institute [43] with a few notable distinctions (fig. 4). At the parcellation level, we see a moderately strong positive correlation between functional activation and structural projection (fig. 4a), which is weaker at the voxel level (fig. 4b). In the midbrain, the coronal slice map shows areas of increased functional activation with respect to structural projection density in the contralateral VTA and the ipsilateral substantia nigra. Coherent clusters of increased activation are also observed in projection areas, most prominently in the ipsilateral and contralateral dorsomedial striatum (fig. 4c). Parcellation-based distributions (figs. 4d and 4e) show this increased activation map encompassing additional areas in the contralateral hemisphere, in particular the contralateral nucleus accumbens, with activity extending into the claustrum. Areas for which structural projections clearly outweigh the functional response are few and dispersed. These small clusters yield only weak negative contrast distributions and are located predominantly in the cerebellum (fig. 4d).

**Figure 4:**
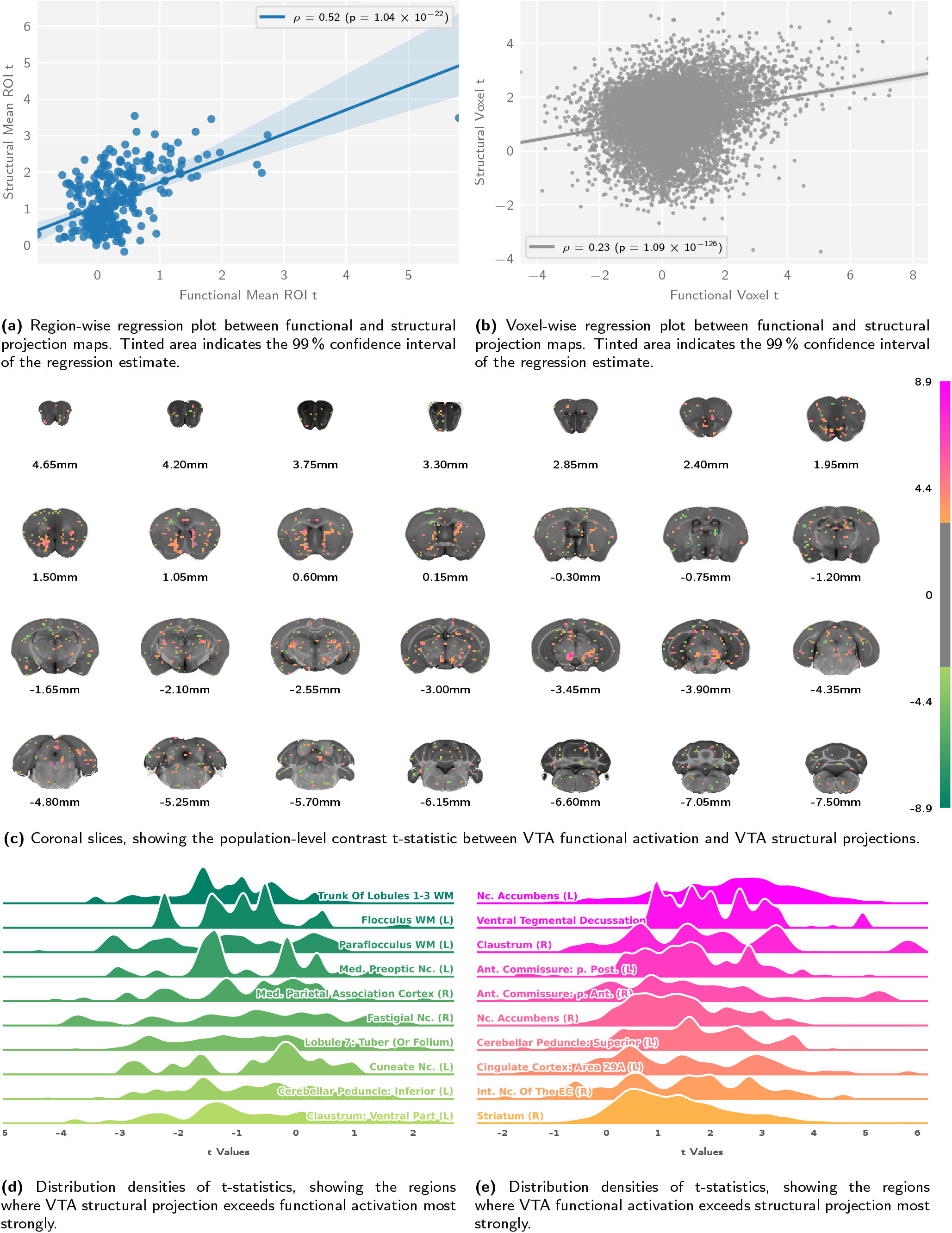
Comparing VTA functional activation to structural projection data reveals good correspondence, with deviations involving the dorsomedial striatum and the contralateral ventral striatum. Depicted are correlation analyses **(a**, **b)** of the population-level functional and structural statistic scores, alongside statistic distributions **(c**, **d**, e) for the contrast, taking into account variability across subjects. Abbreviations: Ant. (Anterior), EC (Endopiriform Claustrum), Int. (Intermediate), Med. (Medial), Nc. (Nucleus), p. (Pars), Post. (Posterior), WM (White Matter),

We differentiate VTA transmission from VTA excitability by mapping functional connectivity using a seed region in the right VTA, which yielded the projection pattern shown in fig. 3e. These clusters are more sparse compared to those identified by stimulus-evoked analysis, yet follow a similar distribution. While areas displaying the highest functional connectivity are located in the right hemisphere, the whole brain parcellation-resolved response displays no significant laterality (*p* = 0.11). Strong activation can be seen in the parcellation regions surrounding the seed, such as the ventral tegmental decussation and the closely located interpeduncular nucleus. In the midbrain, seed-based functional connectivity highlights both the ipsilateral and the contralateral VTA with great specificity, unlike sitmulus-evoked analysis (figs. 3a and 3e). Rostrovental dopaminergic projection areas remain prominently featured, including the nucleus accumbens and the striatum (fig. 3f).

Stimulation in wild type control animals (which is corrected for in the aforementioned stimulus-evoked analyses) does not exhibit a pattern of activity consistent with dopaminergic projections. Sparse grains containing regression scores of *t* ≥ 3 can be observed, with the largest cluster in the lateral geniculate nucleus area of the thalamus, suggesting visual activity (fig. S7b). Atlas parcellation score distributions (fig. S7c) do not strongly deviate from zero, with the highest scoring areas being in the vicinity of the fiber, possibly indicating VTA heating artefacts. Comparable region t-statistic distributions are also found in areas of the cerebellum. Overall the whole brain parcellation-resolved response shows no significant laterality (*p* = 0.68).

Histological analysis of the targeting site reveals that the optic fiber implant displaces the YFP labelled neurons of the VTA (fig. 5). This dislocation was observed irrespective of the targeting area or the speed of implant insertion (10 to 50 μm/s). Yet, labelled filaments and soma remain in the imediate vecinity of the fiber tip, as seen in higher magnification images (fig. 5c).

**Figure 5:**
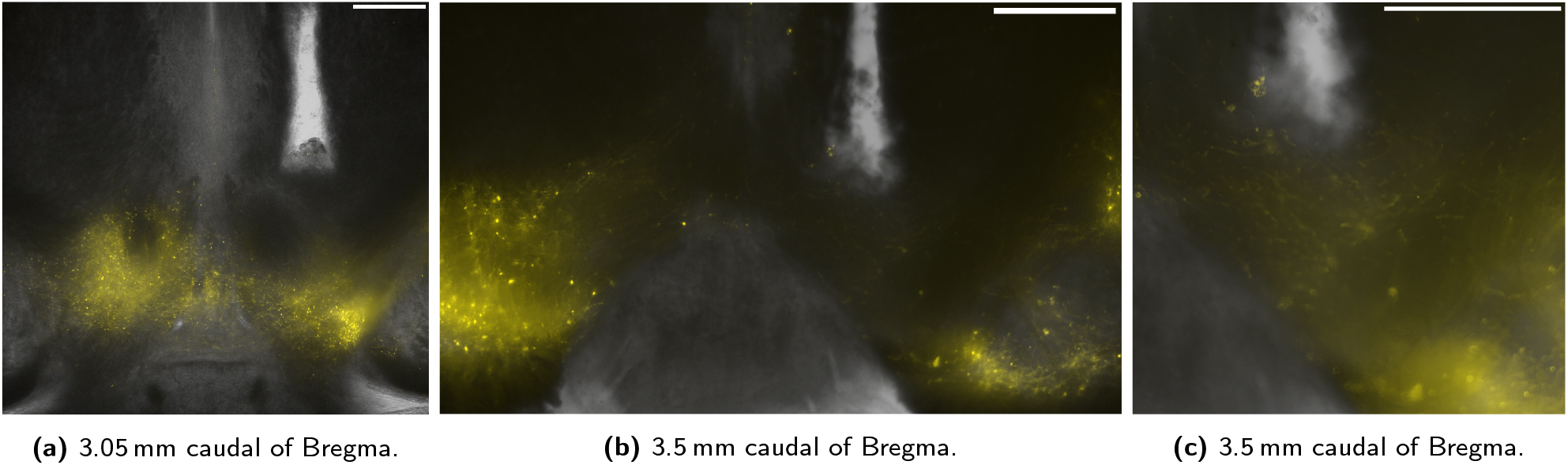
Fiber implantation causes strong local cell displacement in the VTA. Depicted are YFP (coexpressed with Channelrhodopsin) fluorescent microscopy images of the VTA, overlaid on corresponding transmission microscopy images of the same focal plane. All slices are seen in neurological orientation (the right of the image corresponds to the right of the animal). A higher magnification of **(b)** is depicted in **(c)**. White bars indicate a scale of 500 μm, and slices are shown in neurological orientation.

## Discussion

### Whole-Brain Dopaminergic Map

In this article we present the first whole-brain opto-fMRI map of VTA dopaminergic activity in the mouse. Published as voxelwise reusable data and discussed in terms of regions of interest in the article text, this constitutes an essential resource for preclinical investigation of the dopaminergic system. The areas identified as functional VTA dopaminergic targets are largely consistent with histological and electrophysiologic literature (as summarized in fig. 1a). This highlights the suitability of opto-fMRI for interrogating the mouse dopaminergic system, which opens the way for longitudinal recording with whole-brain coverage.

The predominant VTA projection area identified both in literature and in our study is the nucleus accumbens. This area is involved in numerous neuropsychological phenomena, and its activation further supports the method’s suitability to resolve meaningful brain function and increase the predictability of novel interventions using the mouse model organism. Particularly, potential limitations of dopaminergic VTA imaging as shown in recent literature [19], appear to not constrain the protocol detailed in this study.

Throughout brain regions with high signal amplitudes on either metric, we observe a high degree of correspondence between functional activation and structural projection density. Yet, we also document a number of notable differences between opto-fMRI derived projection areas and the structural substrate of the dopaminergic system. Overall, the contrast between function and structure shows stronger signal and wider coverage for the functional activation pattern, particularly in projection areas. Notably, the functional map extends into the contralateral ventral striatum, and both the contralateral and ipsilateral dorsal striatum. Activation of the contralateral ventral striatum might be attributed to an extension of the functional map to the contralateral VTA. This interpretation is supported by the contralateral projection areas showing lower overall significance scores than the ipsilateral areas (figs. 3b and 3f). The explanation of projection area extension into the dorsal striatum on account of secondary activation of the ipsilateral substantia nigra is however less reliable, since the most relevant cluster of increased functional activation — the dorsomedial striatum — can be observed bilaterally, though potential nigral activation is only seen ipsilaterally (fig. 4c). Together with other recent literature [18, 50], it is also possible that VTA activation on its own elicits dorsomedial striatial activity. Not least of all, the local deformation of the VTA upon fiber implantation may additionally confound parcellation in the vicinity of the fiber tip (fig. 5).

Negative contrasts clusters between functional activation and structural projection are overall very sparse (fig. 4d). Yet, the amygdala, hippocampus, and the medial prefrontal cortex — known targets for VTA dopaminergic projections — do not reveal strong activation in opto-fMRI. Comparison with published structural projection data indicates that this is due to low fiber bundle density, as these areas also do not show high amounts of structural projections.

In the pursuit of differentiating primary activation from subsequent signal transmission (and resolving a dopaminergic graph relay model, as depicted in fig. 1b) we present an analysis workflow based on VTA seed-based connectivity. Our results indicate that this analysis is capable of identifying projection areas, but is significantly less powerful than stimulus-evoked analysis (fig. 3a). VTA seed based analysis highlights only a small number of activation clusters and fails to show significant projection laterality. This is an interesting outcome, as — given the superior performance of stimulus-evoked analysis — it describes two possible features of dopaminergic neurotransmission in the VTA. The first is that the relay of primary VTA stimulation has higher fidelity than the fMRI measurement of VTA activity itself (i.e. VTA activity is relayed accurately, but outweighed by measurement noise). The second is that there is a significant threshold to dopaminergic neurotransmission, by which fMRI-measurable baseline activity is predominantly not propagated (i.e. VTA activity is measured accurately, but is relayed in a strongly filtered fashion). The seed-based analysis workflow, however successfully disambiguates VTA activation from adjacent midbrain activation including for the contralateral VTA, which is outside of the seed region of interest. This indicates that VTA susceptibility to optogenetic stimulation may have a unique signature compared to surrounding midbrain tissue in which activation is also elicited in opto-fMRI.

### Assay Parameters

This article presents an evidence-based outline for assay reuse and refinement. In particular, we detail the effects of stimulus protocol categories and optogenetic targeting coordinates on the performance of the method.

The break-down of target coordinates for optical stimulation (fig. 2) indicates that more rostral and deeper implant coordinates elicit stronger VTA signal responses to block stimulation trials. Based on our data we suggest targeting the optic implant at a posteroanterior distance of −3.05 mm from bregma, a left-right distance of 0.5 to 0.55 mm from the midline, and a depth of 4.5 mm from the skull surface. Additional coordinate exploration might be advisable, though further progression towards bregma may lead to direct stimulation of specific efferent fibers rather than the VTA.

The absence of VTA activation as well as coherent activity patterns elicited by phasic stimulation (figs. 2a and S3b) highlights that phasic stimulation is unable to elicit activation measurable by the assay in its current form. The overall low susceptibility to phasic stimulation is most likely due to the intrinsically lower statistical power of such stimulation protocols in fMRI.

Regarding the distribution of activation across projection areas, we note a strong and unexpected divergence between the most sensitive (“best”) and least sensitive (“rejected”) implant coordinate category responses to block stimulation (figs. 3a and 3c). In addition to a difference in VTA and efferent signal intensity (expected as per the selection criterion), we also notice a different pattern of target areas. Interestingly, the activity pattern elicited in the “rejected” group is more strongly weighted towards the hind-brain, and the efferent pattern includes the periaqueductal gray, a prominent brainstem nucleus involved in emotional regulation [51]. This effect might be related to the activation of descending dopaminergic projections, though further investigation is needed to clarify this point and, in general, to better understand the cross-connectivity between deep brain nuclei.

The activation patterns in wild type control animals are very sparse (fig. S7), and — whether or not they are controlled for in the form of a second-level contrast — do not meaningfully impact the dopaminergic block stimulation contrast (figs. 3a and S4). Based on the activation distribution, however, it may be inferred that trace heating artefacts (midbrain activation) and visual stimulation (lateral geniculate nucleus thalamic activation) are present. On account of this, for further experiments, we suggest using eye occlusion and dark or dark-painted ferrule sleeves (to avoid visual stimulation), as well as laser power lower than the 30 mW (239 mW/mm^2^) used in this study (to further reduce heating artefacts).

Stimulus-evoked analysis displayed significant laterality; nevertheless, large clusters displaying significant activation were also observed on the contralateral side. Fluorescence microscopy (fig. 4c) revealed that expression of the viral construct injected at the site of the right VTA extends over a large area, including part of the contralateral VTA. Inspection of the functional map at the midbrain stimulation site corroborates that activity in fact spreads to the contralateral VTA (fig. 3a). This explains the occurrence of contralateral fMRI responses, which are most likely weaker due to a lower photon fluence at the site of the left VTA. Together, these data suggest that the solution volume and virus amount injected for the assay could be significantly reduced, to less than the 0.8 μl (5.7 × 10^12^ vg/ml) used as the minimal volume in this study.

The most salient qualitative feature of fig. 5 is, however, the displacement of labelled neurons from the area in the proximity of the optic fiber implant tip. This feature was consistent across animals and implantation sites, and is a relevant concern as it affects the accuracy of targeting small structures. In particular, such a feature could exacerbate limitations arising from heating artefacts, since the maximum SNR attainable at a particular level of photon fluence may be capped to an unnecessarily low level. This effect might be mitigated by using thinner optic fiber implants (e.g. *ϕ* 200 μm, as opposed to the *ϕ* 400 μm fibers used in this study).

## Conclusion

In this article we demonstrate the suitability of opto-fMRI for investigating a neurotransmitter system which exhibits node-like function in coordinating brain activity. We present the first whole-brain map of VTA dopaminergic signalling in the mouse in a standard space aligned with stereotactic coordinates [46]. We determine that the mapping is consistent with known structural projections, and note the instances where differences are observed. Further, we explore network structure aware analysis via functional connectivity (fig. 3e), finding that the assay provides superior identification of the VTA, but limited support for signal relay imaging. In-depth investigation of experimental variation, based on open source and reusable workflows, supports the current findings by identifying detailed evidence-based instructions for assay reuse. Our study provides a reference dopaminergic stimulus-evoked functional neurophenotype map and a novel and thoroughly documented workflow for the preclinical imaging of dopaminergic function, both of which are crucial to elucidating the etiology of numerous disorders and improving psychopharmacological interventions in health and disease.

## Funding

This work was funded by the Swiss National Science Foundation grant number 310030-179257, which was awarded to MR.

## Supplementary Materials

**Table S1:**
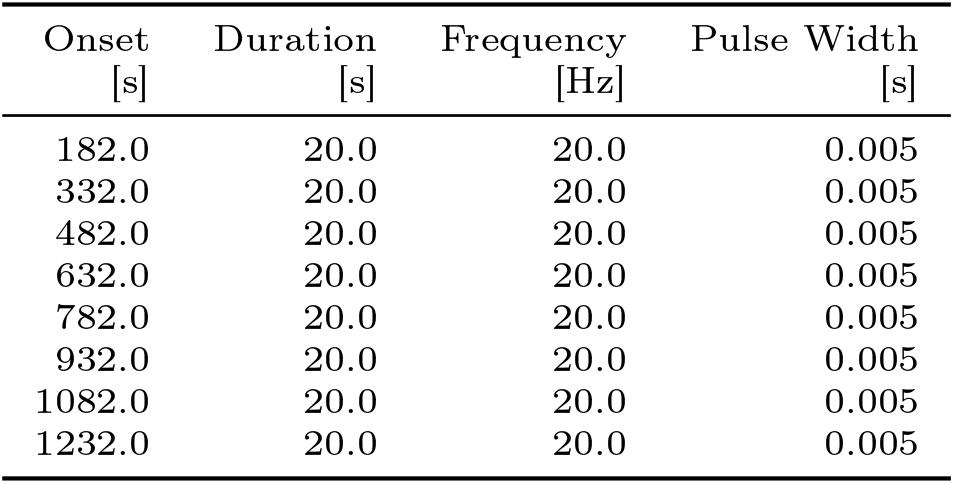
Block stimulation protocol, coded “CogB”.

| Onset [s] | Duration [s] | Frequency [Hz] | Pulse Width [s] |
| --- | --- | --- | --- |
| 182.0 | 20.0 | 20.0 | 0.005 |
| 332.0 | 20.0 | 20.0 | 0.005 |
| 482.0 | 20.0 | 20.0 | 0.005 |
| 632.0 | 20.0 | 20.0 | 0.005 |
| 782.0 | 20.0 | 20.0 | 0.005 |
| 932.0 | 20.0 | 20.0 | 0.005 |
| 1082.0 | 20.0 | 20.0 | 0.005 |
| 1232.0 | 20.0 | 20.0 | 0.005 |

**Table S2:** Block stimulation protocol, coded “CogBr”.

| Onset [s] | Duration [s] | Frequency [Hz] | Pulse Width [s] |
| --- | --- | --- | --- |
| 180.0 | 20.0 | 20.0 | 0.005 |
| 310.0 | 20.0 | 20.0 | 0.005 |
| 480.0 | 20.0 | 20.0 | 0.005 |
| 630.0 | 20.0 | 20.0 | 0.005 |
| 780.0 | 20.0 | 20.0 | 0.005 |
| 930.0 | 20.0 | 20.0 | 0.005 |
| 1080.0 | 20.0 | 20.0 | 0.005 |
| 1230.0 | 20.0 | 20.0 | 0.005 |

**Table S3:** Block stimulation protocol, coded “CogBl”.

| Onset [s] | Duration [s] | Frequency [Hz] | Pulse Width [s] |
| --- | --- | --- | --- |
| 192.0 | 30.0 | 20.0 | 0.005 |
| 342.0 | 30.0 | 20.0 | 0.005 |
| 492.0 | 30.0 | 20.0 | 0.005 |
| 642.0 | 30.0 | 20.0 | 0.005 |
| 792.0 | 30.0 | 20.0 | 0.005 |
| 942.0 | 30.0 | 20.0 | 0.005 |
| 1092.0 | 30.0 | 20.0 | 0.005 |
| 1242.0 | 30.0 | 20.0 | 0.005 |

**Table S4:** Block stimulation protocol, coded “CogBm”.

| Onset [s] | Duration [s] | Frequency [Hz] | Pulse Width [s] |
| --- | --- | --- | --- |
| 180.0 | 8.0 | 20.0 | 0.005 |
| 330.0 | 10.0 | 20.0 | 0.005 |
| 480.0 | 12.0 | 20.0 | 0.005 |
| 630.0 | 14.0 | 20.0 | 0.005 |
| 780.0 | 16.0 | 20.0 | 0.005 |
| 930.0 | 28.0 | 20.0 | 0.005 |
| 1080.0 | 20.0 | 20.0 | 0.005 |
| 1230.0 | 22.0 | 20.0 | 0.005 |

**Table S5:** Block stimulation protocol, coded “CogMwf”.

| Onset [s] | Duration [s] | Frequency [Hz] | Pulse Width [s] |
| --- | --- | --- | --- |
| 150.0000 | 20.0 | 15.0 | 0.005 |
| 280.0000 | 20.0 | 25.0 | 0.005 |
| 410.0000 | 20.0 | 15.0 | 0.010 |
| 540.0000 | 20.0 | 25.0 | 0.010 |
| 670.0000 | 20.0 | 15.0 | 0.005 |
| 799.9999 | 20.0 | 25.0 | 0.005 |
| 930.0000 | 20.0 | 15.0 | 0.010 |
| 1060.0000 | 20.0 | 25.0 | 0.010 |
| 1190.0000 | 20.0 | 15.0 | 0.005 |
| 1320.0000 | 20.0 | 25.0 | 0.005 |
| 1450.0000 | 20.0 | 15.0 | 0.010 |
| 1580.0000 | 20.0 | 25.0 | 0.010 |

**Table S6:** Phasic stimulation protocol, coded “CogP”.

| Onset [s] | Duration [s] | Frequency [Hz] | Pulse Width [s] |
| --- | --- | --- | --- |
| 190.0 | 0.8 | 25.0 | 0.005 |
| 192.0 | 0.8 | 25.0 | 0.005 |
| 194.0 | 0.8 | 25.0 | 0.005 |
| 196.0 | 0.8 | 25.0 | 0.005 |
| 290.0 | 0.8 | 25.0 | 0.005 |
| 292.0 | 0.8 | 25.0 | 0.005 |
| 294.0 | 0.8 | 25.0 | 0.005 |
| 296.0 | 0.8 | 25.0 | 0.005 |
| 390.0 | 0.8 | 25.0 | 0.005 |
| 392.0 | 0.8 | 25.0 | 0.005 |
| 394.0 | 0.8 | 25.0 | 0.005 |
| 396.0 | 0.8 | 25.0 | 0.005 |
| 490.0 | 0.8 | 25.0 | 0.005 |
| 492.0 | 0.8 | 25.0 | 0.005 |
| 494.0 | 0.8 | 25.0 | 0.005 |
| 496.0 | 0.8 | 25.0 | 0.005 |
| 590.0 | 0.8 | 25.0 | 0.005 |
| 592.0 | 0.8 | 25.0 | 0.005 |
| 594.0 | 0.8 | 25.0 | 0.005 |
| 596.0 | 0.8 | 25.0 | 0.005 |

**Table S7:**
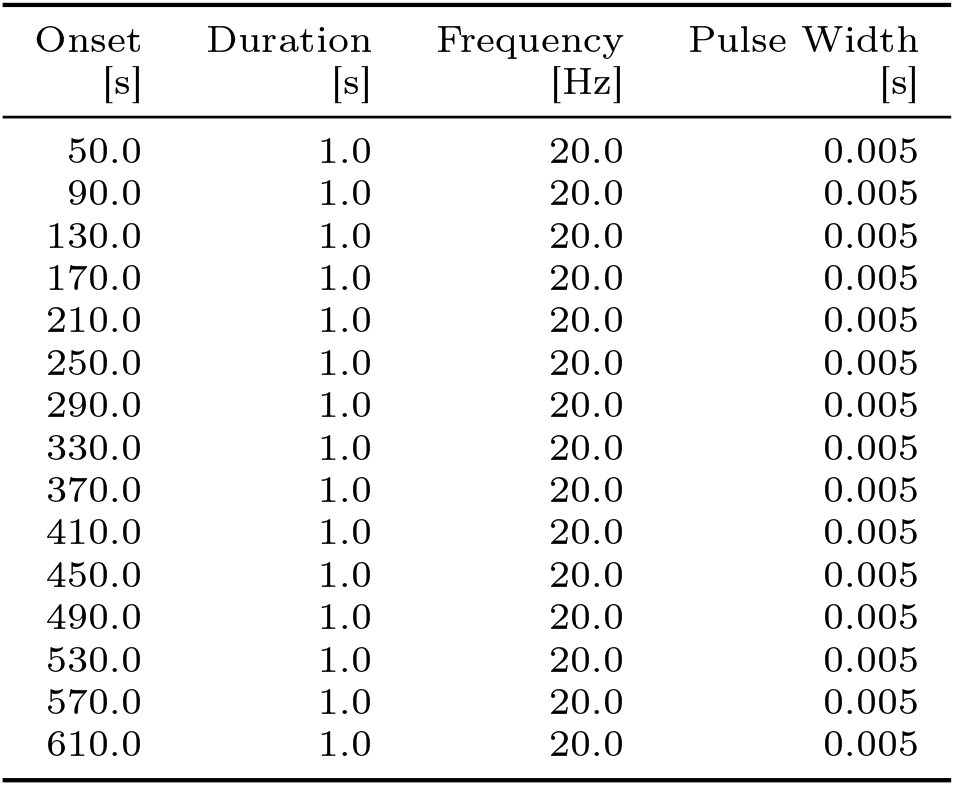
Phasic stimulation protocol, coded “JPogP”.

| Onset [s] | Duration [s] | Frequency [Hz] | Pulse Width [s] |
| --- | --- | --- | --- |
| 50.0 | 1.0 | 20.0 | 0.005 |
| 90.0 | 1.0 | 20.0 | 0.005 |
| 130.0 | 1.0 | 20.0 | 0.005 |
| 170.0 | 1.0 | 20.0 | 0.005 |
| 210.0 | 1.0 | 20.0 | 0.005 |
| 250.0 | 1.0 | 20.0 | 0.005 |
| 290.0 | 1.0 | 20.0 | 0.005 |
| 330.0 | 1.0 | 20.0 | 0.005 |
| 370.0 | 1.0 | 20.0 | 0.005 |
| 410.0 | 1.0 | 20.0 | 0.005 |
| 450.0 | 1.0 | 20.0 | 0.005 |
| 490.0 | 1.0 | 20.0 | 0.005 |
| 530.0 | 1.0 | 20.0 | 0.005 |
| 570.0 | 1.0 | 20.0 | 0.005 |
| 610.0 | 1.0 | 20.0 | 0.005 |

In a linear modelling of the implant coordinate variables, the VTA mean t statistic is found sensitive only to the stimulation protocol category (*F*_1,59_ = 57.3, *p* = 2.92 × 10^−10^), but not the stimulation target depth (*F*_1,59_ = 0.48, *p* = 0.49), the stimulation target posteroanterior (PA) coordinates (*F*_1,59_ = 0.59, *p* = 0.45), and the interaction of the depth and PA target coordinates (*F*_1,59_ = 0.48, *p* = 0.49).

**Figure S1:**
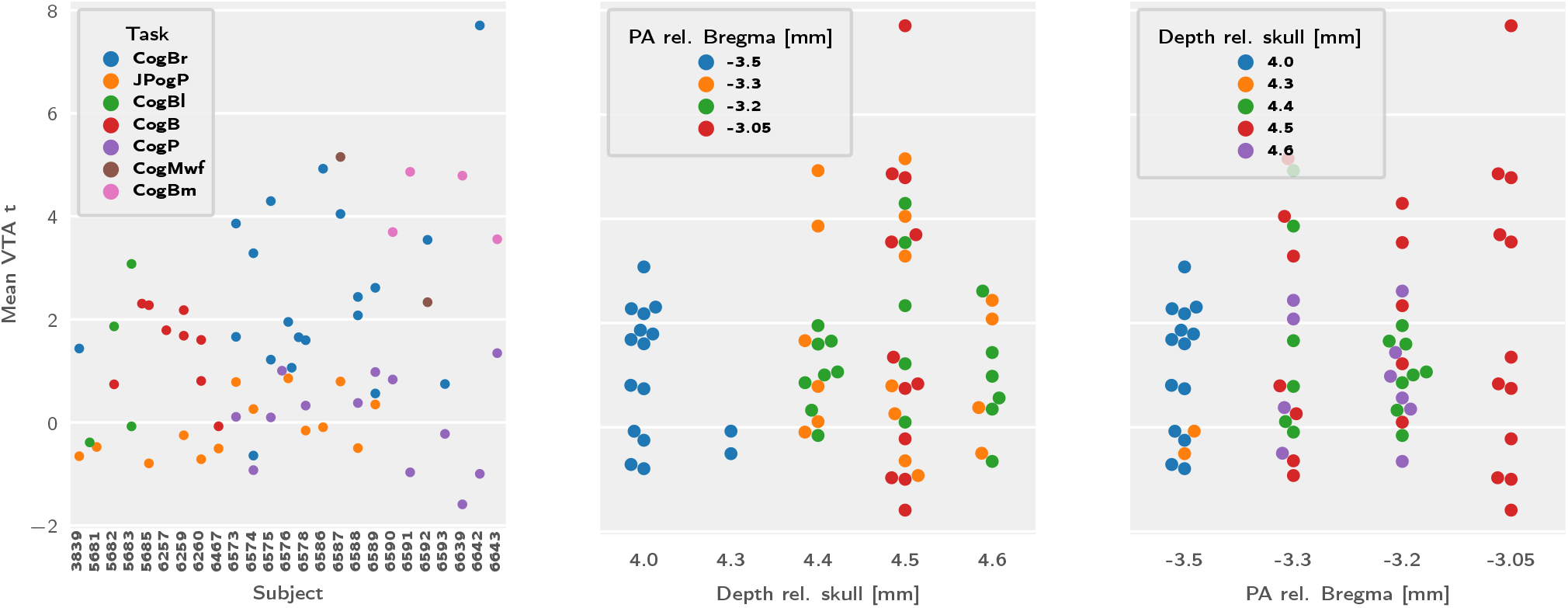
Multifactorial (depth and posteroantior) implant coordinate comparisons of signal intensity in the VTA region of interest. Protocols coded as in tables S1 to S7.

**Figure S2:**
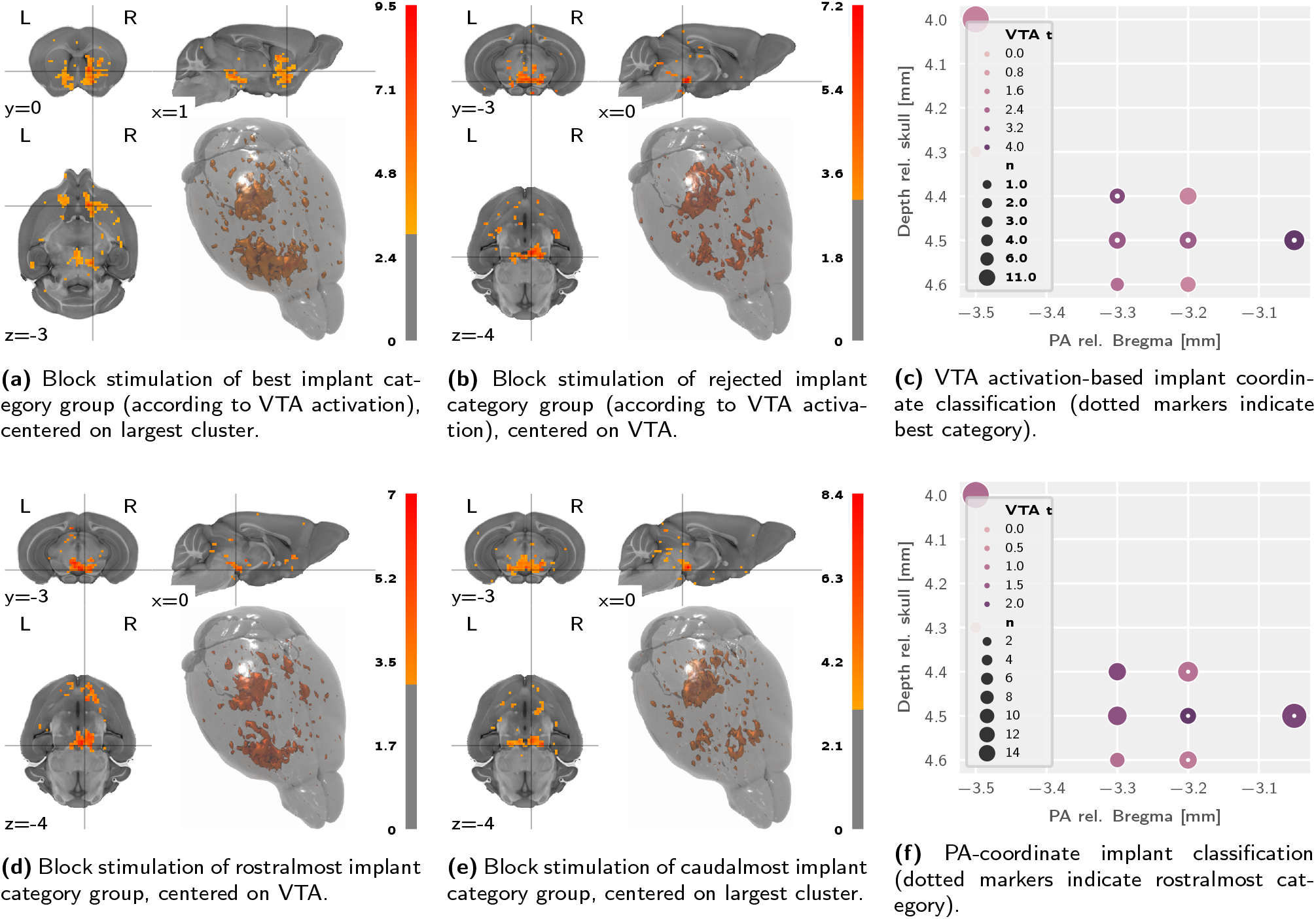
PA-coordinate-based classification does not show a better projection segmentation than block trial-based classification. Depicted are t-statistic maps (centerd on largest cluster, thresholded at t ≥ 3) of the second-level analysis for block stimulation protocols, divided into best and rejected **(a**, **b)**, or rostralmost and caudalmost **(d**, **e)**. All maps are adjusted for the wild type control stimulation effects.

**Figure S3:**
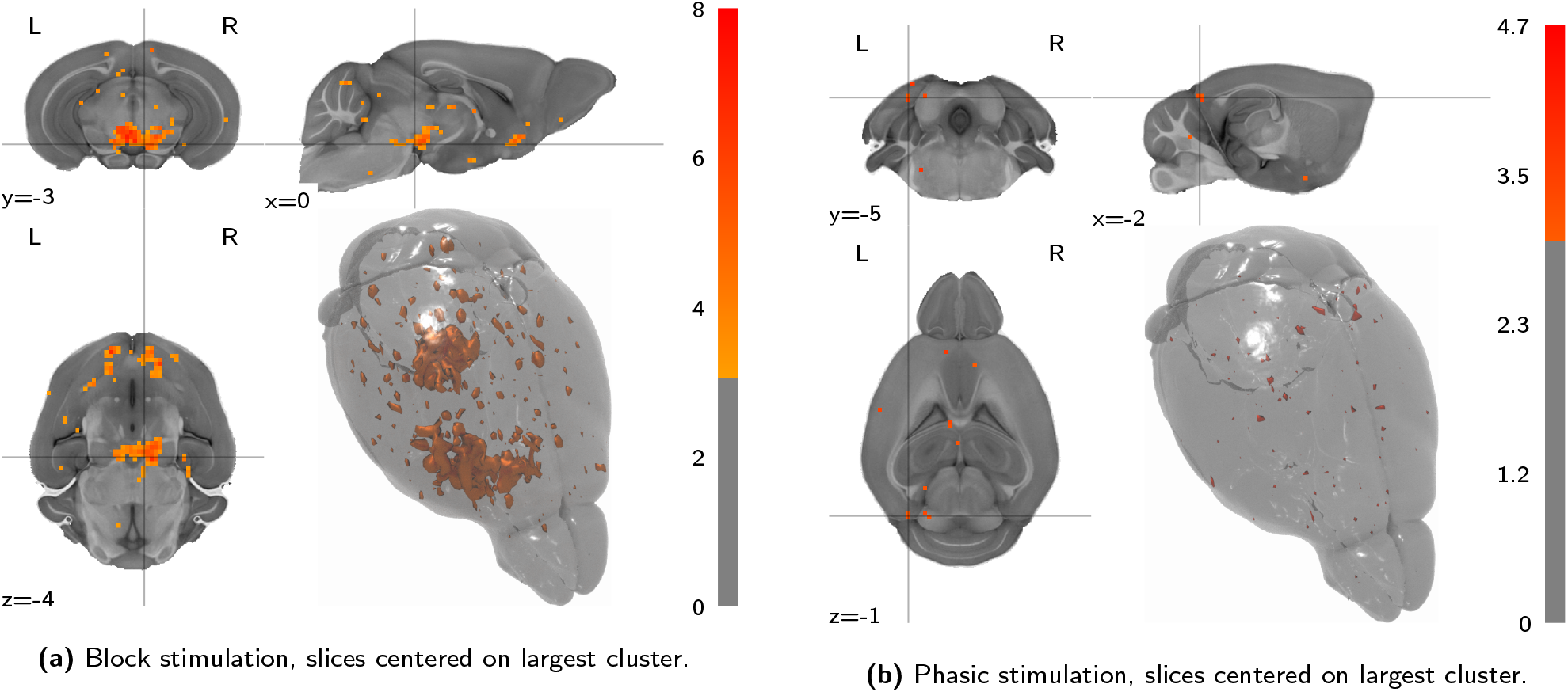
No negative activation patterns are salient upon block VTA stimulation, and no coherent activation patterns of any sort after phasic VTA stimulation. Depicted are t-statistic maps (thresholded at |t| ≥ 3) of second-level analyses, divided by stimulation category and binning all implant coordinates. Slices are centered on the VTA coordinates (RAS = 0.5/ − 3.2/ − 4.5) and the largest cluster, respectively. All maps are adjusted for the wild type control stimulation effects.

**Figure S4:**
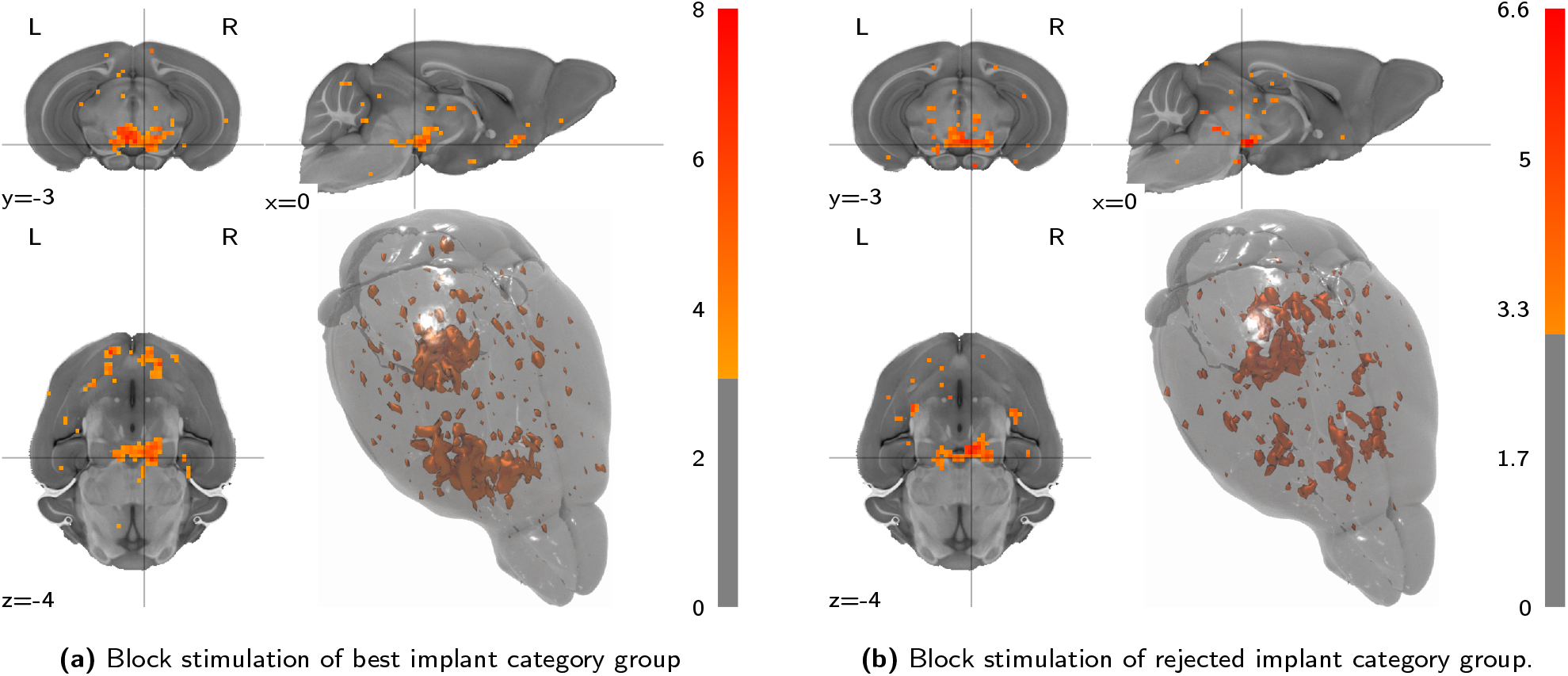
The uncorrected population-level response to block stimulation does not qualitatively differ from the wild type control corrected results in figs. 3a and 3c. Depicted are wildtype-control-uncorrected t-statistic maps (thresholded at t ≥ 3) of the second-level analysis for block stimulation protocols, divided by implant category group. Slices are centered on the VTA region of interest.

**Figure S5:**
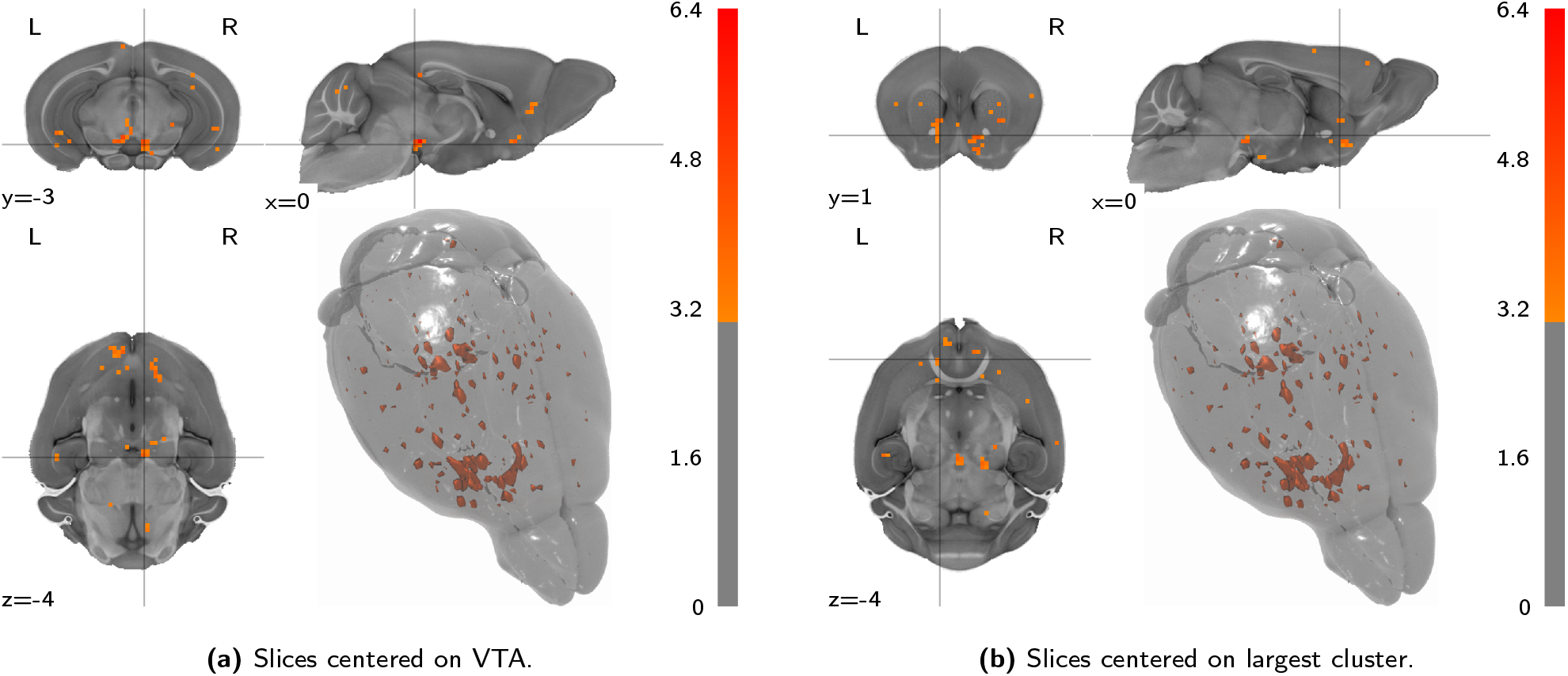
Depicted are t-statistic maps (thresholded at t ≥ 3) of the second-level analysis for block stimulation task VTA seed functional connectivity, observed in the best implant category, corrected for the negative control baseline. Slices are centered on the VTA coordinates (RAS = 0.5/ − 3.2/ − 4.5) and the largest cluster, respectively. This comparison is only provided for the sake of completeness and analogy with the stimulus-evoked analysis. Conceptually this comparison is not of primary interest, since seed-based functional connectivity attempts to include precisely the baseline functioning of the system into the evaluation.

**Figure S6:**
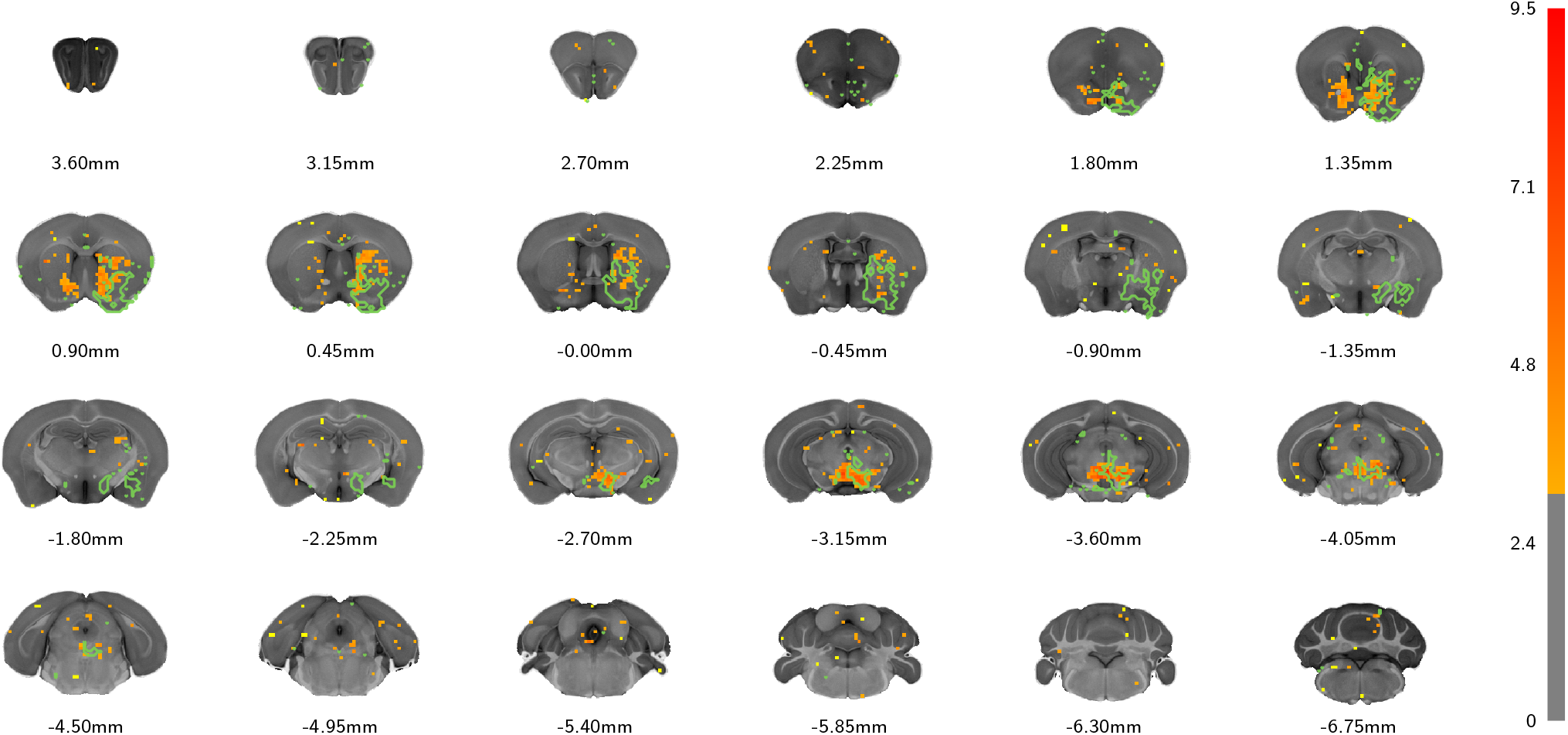
Coronal slice overlay, showing the VTA functional activation t-statistic heatmap (as in **fig. 3a**), and the VTA structural projection outline, both thresholded at t ≥ 3. Interpretation of this figure as showcasing a complementarity in the patterns is cautioned, as qualitative inspection of thresholded data does not accurately capture variation in the statistic distributions. For statements regarding the compariosn of functional activation and structural projection, figs. 4a to 4c are more suitable.

**Figure S7:**
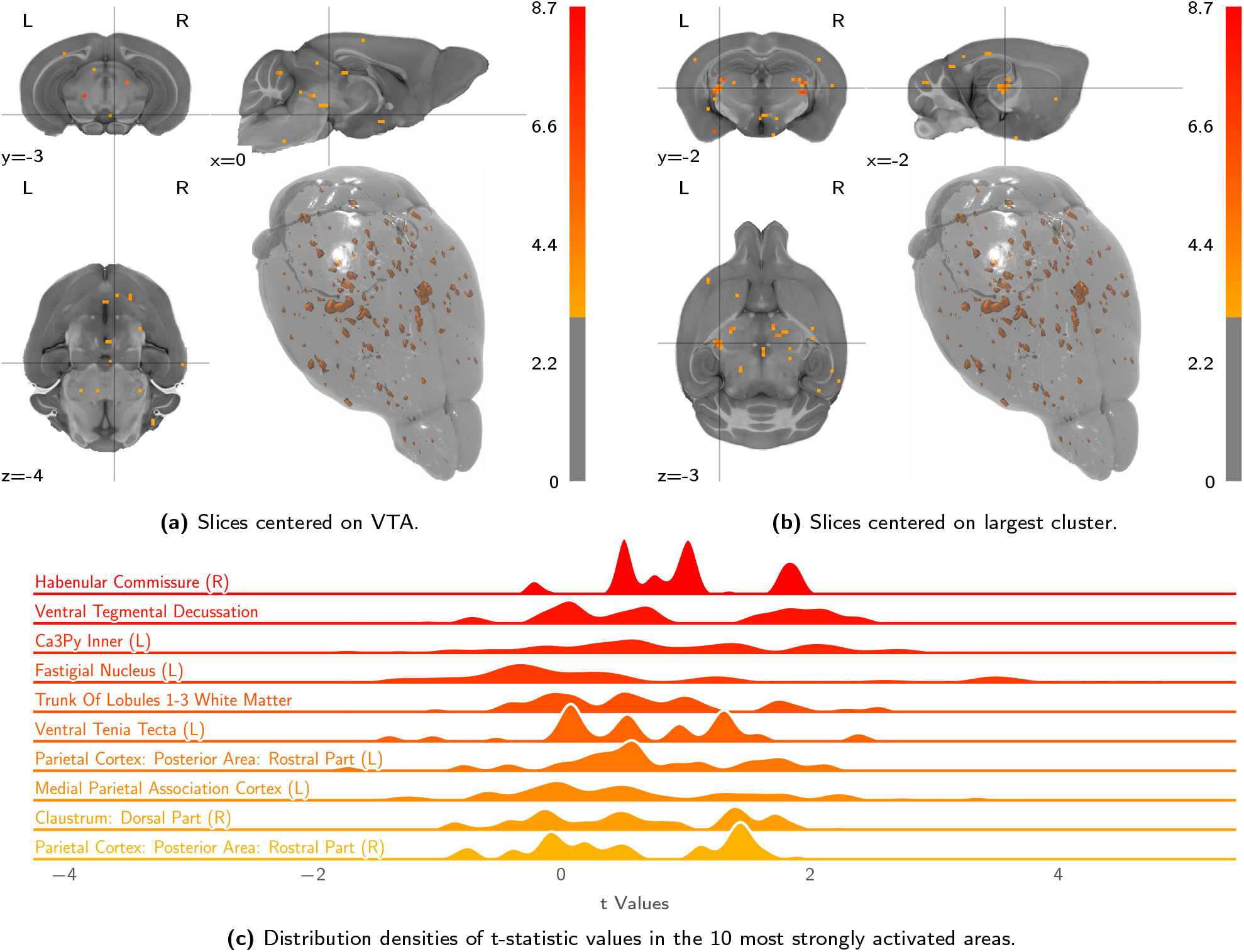
Block stimulation in wild type control animals produces no large activation clusters, yet scattered activation hints at some visual excitation and heating artefacts. Depicted are volumetric population t-statistic maps **(a**, **b)** — thresholded at t ≥ 3, as well as a break-down of activation along atlas parcellation regions **(c)**.

## Notes

https://zenodo.org/record/3575149/files/opfvta_bidsdata-2.0.tar.xz

### Competing Interest Statement

The authors have declared no competing interest.

### Summary of Updates

formatting and background section improvement. Corrected source data link.

